# Genetic ancestry and population structure in the All of Us Research Program cohort

**DOI:** 10.1101/2024.12.21.629909

**Authors:** Shivam Sharma, Shashwat Deepali Nagar, Priscilla Pemu, Stephan Zuchner, SEEC Consortium, Leonardo Mariño-Ramírez, Robert Meller, I. King Jordan

## Abstract

The NIH All of Us Research Program (*All of Us*) aims to build one of the world’s most diverse population biomedical datasets in support of equitable precision medicine. For this study, we analyzed participant genomic variant data to assess the extent of population structure and to characterize patterns of genetic ancestry for the *All of Us* cohort (n=297,549). Unsupervised clustering of genomic principal component analysis (PCA) data revealed a non-uniform distribution of genetic diversity and substantial population structure in the *All of Us* cohort, with dense clusters of closely related participants interspersed among less dense regions of genomic PC space. Supervised genetic ancestry inference was performed using genetic similarity between *All of Us* participants and global reference population samples. Participants show diverse genetic ancestry, with major contributions from European (66.4%), African (19.5%), Asian (7.6%), and American (6.3%) continental ancestry components. Participant genetic similarity clusters show group-specific genetic ancestry patterns, with distinct patterns of continental and subcontinental ancestry among groups. We also explored how genetic ancestry changes over space and time in the United States (US). African and American ancestry are enriched in the southeast and southwest regions of the country, respectively, whereas European ancestry is more evenly distributed across the US. The diversity of *All of Us* participants’ genetic ancestry is negatively correlated with age; younger participants show higher levels of genetic admixture compared to older participants. Our results underscore the ancestral genetic diversity of the *All of Us* cohort, a crucial prerequisite for genomic health equity.

## Introduction

The biomedical research community has become increasingly aware of the genomics research gap, whereby the vast majority of participants in genetics research cohorts are of European ancestry^1, 2, 3^. The Eurocentric bias in genomics research threatens to exacerbate health disparities, since discoveries made with European ancestry cohorts may not transfer to diverse ancestry groups^4^. The NIH All of Us Research Program (*All of Us*) is a large cohort study of people who live in the US that combines participant genomic, phenotypic, and environmental data, with health-related outcome data gleaned from surveys and electronic health records^5, 6^. *All of Us* has emphasized the recruitment of participants from population groups that are underrepresented in biomedical research in an effort to close the genomics research gap and to ensure that the benefits of precision medicine are shared equitably among all people^7, 8^.

*All of Us* demonstration projects are being used to describe and validate the initial genomic data release and the cloud-based Researcher Workbench, where registered users can access and analyze participant data^9^. The aim of this demonstration project was to characterize the patterns of population structure and genetic ancestry among *All of Us* participants. Population structure refers to differences in the frequencies of genetic variants (alleles) among different groups or populations within a species, and population structure can be revealed by the presence of clusters of genetically similar individuals^10^. Genetic ancestry is closely related to the concept of population structure, and it can be defined conceptually, mechanistically, and operationally. Conceptually, genetic ancestry reflects the geographic origins of an individual’s ancestors^11, 12, 13, 14^. Mechanistically, genetic ancestry has been defined as the subset of genealogical paths through which an individual’s DNA has been inherited from their ancestors^15^. For any individual, only a subset of their genealogical ancestors contributes DNA to their genome. Operationally, genetic ancestry is typically characterized via genetic similarity between query individuals (e.g. *All of Us* participants) and individuals from global reference populations, which are taken as surrogates for ancestral populations^16, 17, 18, 19^.

For this demonstration study of the *All of Us* cohort, we analyzed participant genomic variant data to (1) assess the extent of population structure in the cohort, (2) characterize the patterns of participant genetic ancestry at continental and subcontinental levels, and (3) explore how participants’ genetic ancestry changes over space and time in the US. Our results reveal substantial population structure and heterogeneous patterns of genetic ancestry among *All of Us* participants, consistent with the consortium’s efforts to recruit a diverse participant cohort.

## Materials and Methods

### All of Us participant cohort, consent, and IRB review

This study was performed as an *All of Us* genomic data demonstration project^5^. *All of Us* demonstration projects are intended to describe and validate data and analysis tools for the participant cohort. Details on the initial *All of Us* data release and Researcher Workbench used for this study were previously published^6^. The genomic data demonstration project and experimental protocols were approved by the *All of Us* Institutional Review Board (#2016–05-TN-Master), and informed consent was obtained from all participants. *All of Us* inclusion criteria include adults 18 and older, with the legal authority and decisional capacity to consent, and currently residing in the US or a territory of the US. *All of Us* exclusion criteria exclude minors under the age of 18 and vulnerable populations (prisoners and individuals without the capacity to give consent). Details on participant recruitment, informed consent, inclusion and exclusion criteria are available online at https://allofus.nih.gov/sites/default/files/All_of_Us_operational_protocol_v1.7_mar_2018.pdf. Results reported here comply with the *All of Us* Data and Statistics Dissemination Policy disallowing disclosure of group counts under 20.

The *All of Us* Researcher Workbench was used to build the participant cohort for this study (Supplementary Figure 1). The cohort was built from the *All of Us* Controlled Tier dataset v7 (curated version C2022Q4R9), which includes participants enrolled from 2018-2022, with a data cutoff date of 7/1/2022. Participants who self-identified as American Indian or Alaska Native were not included in the analysis.

### Unsupervised genetic clustering analysis

Participant genomic data were accessed from the Controlled Tier dataset. Genome-wide genotypes for *All of Us* participants were characterized using the Illumina Global Diversity Array with variants called for 1,824,517 genomic positions on the GRCh38/hg38 reference genome build. *All of Us* participant variants were merged and harmonized with whole genome sequence variant data from 3,433 global reference samples characterized as part of the 1000 Genomes Project (1KGP; phase 3) and the Human Genome Diversity Project (HGDP; Supplementary Table 1)^20, 21^. Biallelic variants common to the *All of Us* and reference data sets were merged, with strand flips and variant identifier inconsistencies harmonized as needed. Variants with >1% missingness and <1% minor allele frequency were removed from the merged and harmonized dataset. Linkage disequilibrium (LD) pruning was done using window size=50, step size=10, and pairwise threshold r^2^<0.1, yielding a final *All of Us* and global reference sample dataset of 187,795 variants. Variant merging, harmonization, and LD pruning were performed using PLINK version 1.9^22^ and custom scripts as previously described^23, 24, 25^. The final dataset of *All of Us* participant genomic variants was used for unsupervised clustering analysis. Principal Component Analysis (PCA) was run on the variant dataset using the FastPCA program implemented in PLINK version 2.0. The clustering tendency of the resulting genomic PCA data was analyzed using the Hopkins statistic with the hopkins R package^26^ and nearest neighbor search with the FNN R package version 1.1.4^27^. Kernel density estimation was performed with the MASS R package using PCs 1-3 and contour lines were extracted from the estimated density distribution^28^. Density-based clustering was performed using the HDBSCAN algorithm^29^. HDBSCAN was run on first 5 PCs for the PCA data with parameters min_samples=2,000 and min_cluster_size=2,500. Cluster boundaries were visualized using the ggforce R package.

### Supervised genetic ancestry inference

Genomic variants from *All of Us* participants and a set of four global reference populations were merged and harmonized as described in the previous section to perform continental and subcontinental genetic ancestry inference. Kinship analysis was performed with the KING program to eliminate related (or duplicated) reference samples from the global reference populations^30^. Continental genetic ancestry inference was performed using a subset of 1,572 global reference samples from the 1KGP and the HGDP, which were selected as non-admixed representatives of seven ancestry groups: African, American, East Asian, South Asian, West Asian, European, and Oceanian (Supplementary Table 1). K-nearest neighbor clustering of genomic PCA data was used to identify *All of Us* participants that cluster together with African, East Asian, South Asian, and European reference populations, and these participants were used for subcontinental ancestry inference^31^. West Asian and Oceanian reference populations were not used for this purpose owing to the relatively low number of participants that clustered with these groups. Asian and European reference populations for subcontinental ancestry inference were taken from the 1KGP and HGDP (Supplementary Table 2). 1KGP and HGDP reference populations were used together with additional reference populations to provide broader geographic coverage for African and American subcontinental ancestry inference (Supplementary Table 2). African reference samples were taken from a study of Bantu-speaking populations in Africa that included samples from 53 populations from east, central, south, and west Africa^32^. The merged and harmonized African subcontinental ancestry inference panel included 1,659 reference samples and 228,033 variants.

Continental and subcontinental ancestry inference was performed via analysis of merged *All of Us* participant and global reference population genomic variant sets with the program Rye (<u>R</u>apid ancestr<u>Y</u> <u>E</u>stimation)^33^. Rye performs rapid and accurate genetic ancestry inference based on principal component analysis (PCA) of genomic variant data. PCA was run on the merged variant datasets using the FastPCA program implemented in PLINK version 2.0, and Rye was then run on the first 25 PCs, using the defined reference ancestry groups to assign ancestry group fractions to individual *All of Us* participant samples. The continuous ancestry fractions that we report here were calculated independently of the categorical ancestry predictions currently provided by the *All of Us* Researcher Workbench^34^.

*All of Us* participant continental ancestry fractions were visualized as admixture-style plots at the state (or territory) level using the geofacets R package^35, 36^. Admixture entropy (*AE*) was used to quantify the amount of genetic admixture for *All of Us* participants as previously described^25, 37^: 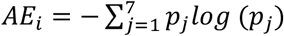, where *p”* is the fraction of ancestry group *j* for individual *i*.

### Note on genetic ancestry inference

As discussed in the introduction, genetic ancestry can be defined conceptually, mechanistically, and operationally. We use an operational definition of genetic ancestry for *All of Us* participants in this study, as measured by their levels of genetic similarity with global reference population samples^16, 17^. Accordingly, the phrase ‘African ancestry’ is used here as shorthand for similarity to African reference population samples, ‘European ancestry’ is used for similarity to European reference population samples and so on. ‘American ancestry’ refers to genetic similarity in Indigenous American reference population samples. The relative levels of similarity to different reference population groups allows us to infer percent ancestry components for *All of Us* participants^33^. The genetic ancestry results reported here are contingent upon the choice of reference populations, how these reference populations are delineated, and the method used to infer genetic similarity between *All of Us* participants and the reference population samples.

## Results

### Unsupervised: population structure

A cohort of 297,549 *All of Us* participants, for whom genomic data are available, was created using the *All of Us* Researcher Workbench (Supplementary Figure 1). *All of Us* participant genetic diversity was analyzed using PCA of genomic variant data followed by unsupervised clustering to assess the extent of population structure in the cohort. The clustering tendency of participant genomic PCA data was evaluated using the Hopkins statistic, nearest neighbors, and kernel density estimation. The PCA data yield a Hopkins statistic value of ∼1, indicating highly clustered, non-uniformly, and non-randomly distributed genomic PCA data. The numbers of close neighbors per participant are highly variable across PC space, and kernel density estimation shows a multimodal distribution with distinct peaks separated in PC space (Figure 1A and 1B). All three of these metrics reveal highly clustered participant genomic data, with dense groups of genetically similar individuals interspersed among less dense regions, indicative of substantial population structure in the *All of Us* cohort.

**Figure 1.**
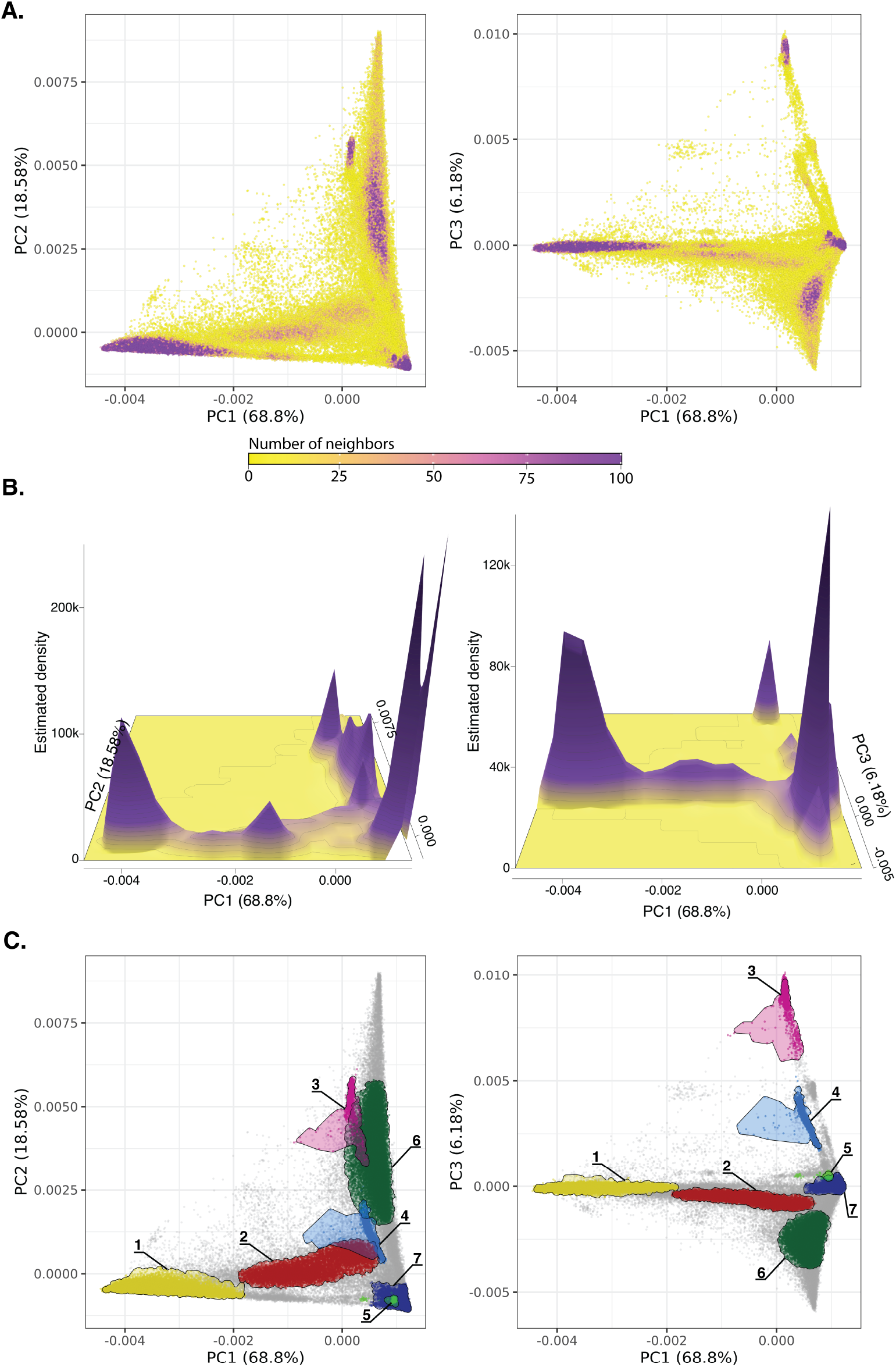
Population structure. Genomic PCA for *All of Us* participants. Left panels show PC1 versus PC2 comparisons, and right panels show PC1 versus PC3 comparisons, with the percent of variance explained by each PC shown. (A) Participants color-coded by the number of close neighbors as defined by Euclidean distance<0.1 in PCs 1-5. (B) Kernel density estimation with peaks showing high density clusters of participants in PC space. (C) High density clusters of genetically similar participants shown as groups 1-7.

Density-based clustering of the genomic PCA data yield an optimal number of K=7 genetic diversity clusters (Figure 1C). Similar clustering was performed using a Uniform Manifold Approximation and Projection (UMAP) analysis of the genomic PCA data (Supplementary Methods). Density-based clustering of UMAP data reveals almost twice as many clusters (K=13) as seen for the PCA data, but there is broad concordance between the two methods with high percentages of participant overlap for each PCA cluster within one or two corresponding UMAP clusters (Supplementary Figure 2). The number of *All of Us* genetic diversity clusters could change with future participant data releases.

### Supervised: genetic ancestry

*All of Us* participant genetic ancestry was inferred using genomic PCA data analyzed with the Rye (<u>R</u>apid ancestr <u>Y</u> <u>E</u>stimation) program^33^. Participant PCA data were compared with PCA data from global reference populations, taken from the 1KGP and the HGDP, to infer individual ancestry proportions from seven continental-level ancestry groups: African, American, East Asian, South Asian, West Asian, European, and Oceanian (Supplementary Table 1 and Supplementary Figure 3). *All of Us* participants are broadly distributed in PC space, whereas global reference samples from different ancestry groups are tightly clustered in PC space (Figure 2A and 2B). Rye infers *All of Us* participant genetic ancestry proportions as linear combinations of reference population ancestries. Overall, the *All of Us* participant cohort shows 19.51% African, 6.33% American, 2.57% East Asian, 3.05% South Asian, 1.95% West Asian, 66.37% European, and 0.21% Oceanian ancestry. The *All of Us* participant genetic similarity groups inferred with density-based clustering show group-specific patterns of ancestry proportions, with a continuum of ancestry proportions within and between groups (Figure 2C). Groups 1, 3, 4, and 7 show the most uniform patterns of ancestry within groups, whereas groups 2, 5, 6, and the remaining participants that did not fall into any density-based cluster show more diverse patterns of ancestry and admixture. All groups show evidence of admixture with multiple ancestry components present in different proportions.

**Figure 2.**
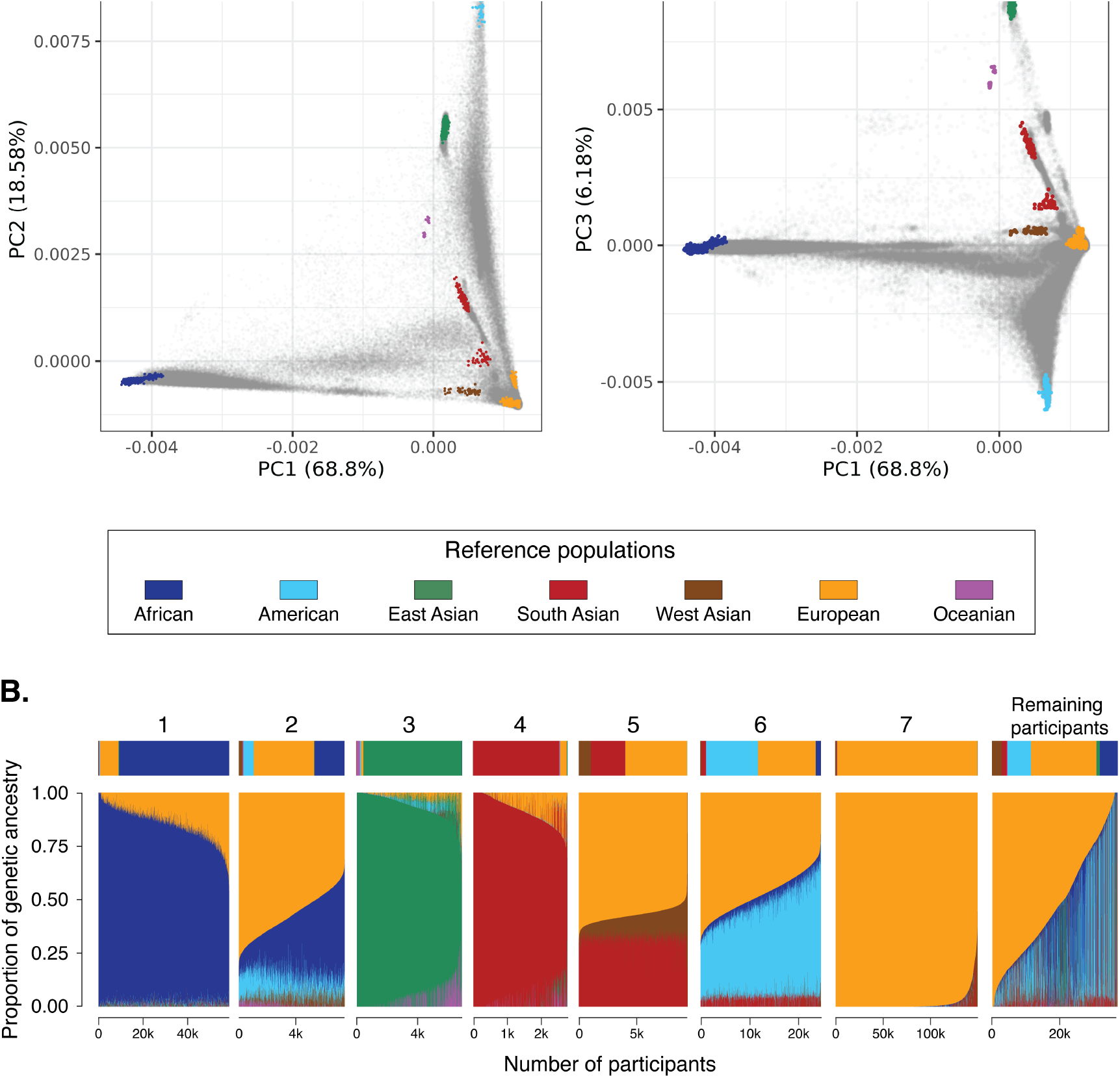
Continental genetic ancestry. (A) Genomic PCA with *All of Us* participants shown in gray and global reference population samples color-coded as shown in the key. Left panels show PC1 versus PC2 comparisons, and right panels show PC1 versus PC3 comparisons, with the percent of variance explained by each PC shown. (B) Genetic ancestry proportions for *All of Us* participants stratified by the genetic similarity groups shown in Figure 1C. Average ancestry proportions are shown above each group, and numbers of participants are shown below each group. The remaining participants are individuals that did not fall into a dense PCA cluster.

The *All of Us* Researcher Workbench predicts participant membership among six continental ancestry groups, using a PCA-based machine learning method that is distinct from the continuous ancestry inference approach used here^34^. We compared the participant continental ancestry percentages inferred here to the Researcher Workbench assigned categorical ancestry groups (Supplementary Figure 4). Five of the six categorical ancestry groups correspond exactly with the reference population groups we use: African, East Asian, South Asian, Middle Eastern (West Asian here), and European. For these five groups, there is high correspondence between participants’ PCA-based machine learning predicted group membership and averages for the ancestry percentages that we inferred (83.02-97.71% matching ancestry). The Admixed American ancestry category from the Researcher Workbench includes modern, admixed reference samples from Latin America, whereas our American reference population group includes Indigenous American samples only (Supplementary Table 1). The Admixed American group shows 51.01% European ancestry and 35.84% American ancestry, consistent with what is expected for modern Latin American populations^38, 39^.

We also used Rye to infer subcontinental ancestry for *All of Us* participants with high levels of African (n=9,291), East Asian (n=2,457), South Asian (n=2,484), and European ancestry (n=24,730; Figure 3). The relationships among the reference populations used for subcontinental ancestry inference with Rye and *All of Us* participants are shown in Supplementary Figures 5-7. African subcontinental ancestry is characterized by a predominant West Central African component, followed by West African and Bantu components. East Asian subcontinental ancestry is highly diverse with predominant Han (Chinese), Japanese, and Southeast Asian components. South Asian subcontinental ancestry is mainly South Indian followed by North Indian and a small Central Asian component. European subcontinental ancestry is made up primarily of British ancestry followed by Italian and Iberian components.

**Figure 3.**
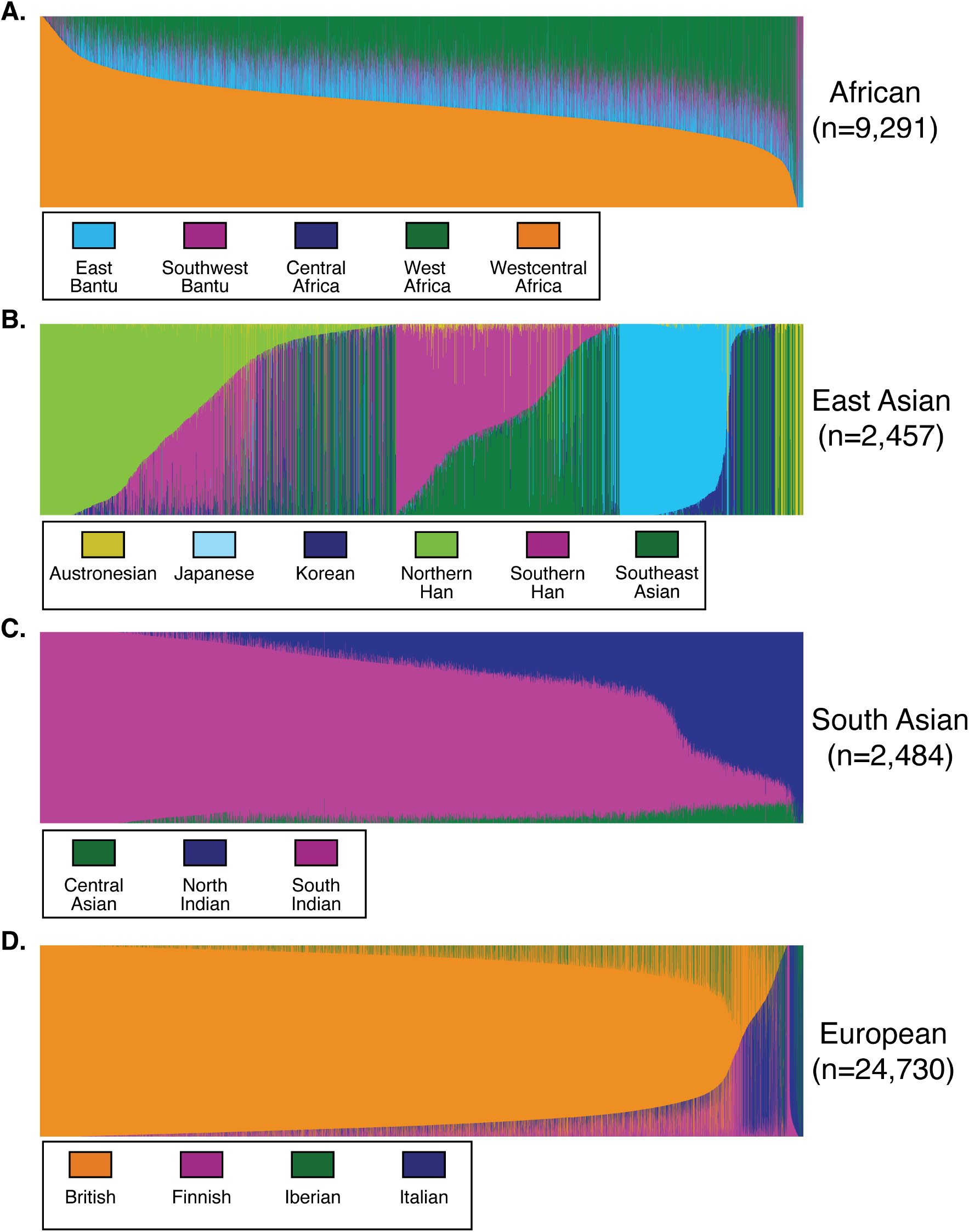
Subcontinental genetic ancestry. Subcontinental genetic ancestry proportions for *All of Us* participants from African, American, East Asian, South Asian, and European continental ancestry groups. Subcontinental groups (regions) for each continental ancestry group are color-coded as shown.

### Genetic ancestry by geography and age

*All of Us* participant continental ancestry percentages were visualized across fifty states and Puerto Rico to evaluate the geographic distribution of ancestry across the US (Figure 4). African ancestry is concentrated primarily in the southeast part of the country, whereas American ancestry if found primarily in the southwest and California. European ancestry is more uniformly distributed across the country, with the highest concentrations found in north, along the Canadian border. Relatively high levels of admixture are seen in the northeast, Florida, and Hawaii.

**Figure 4.**
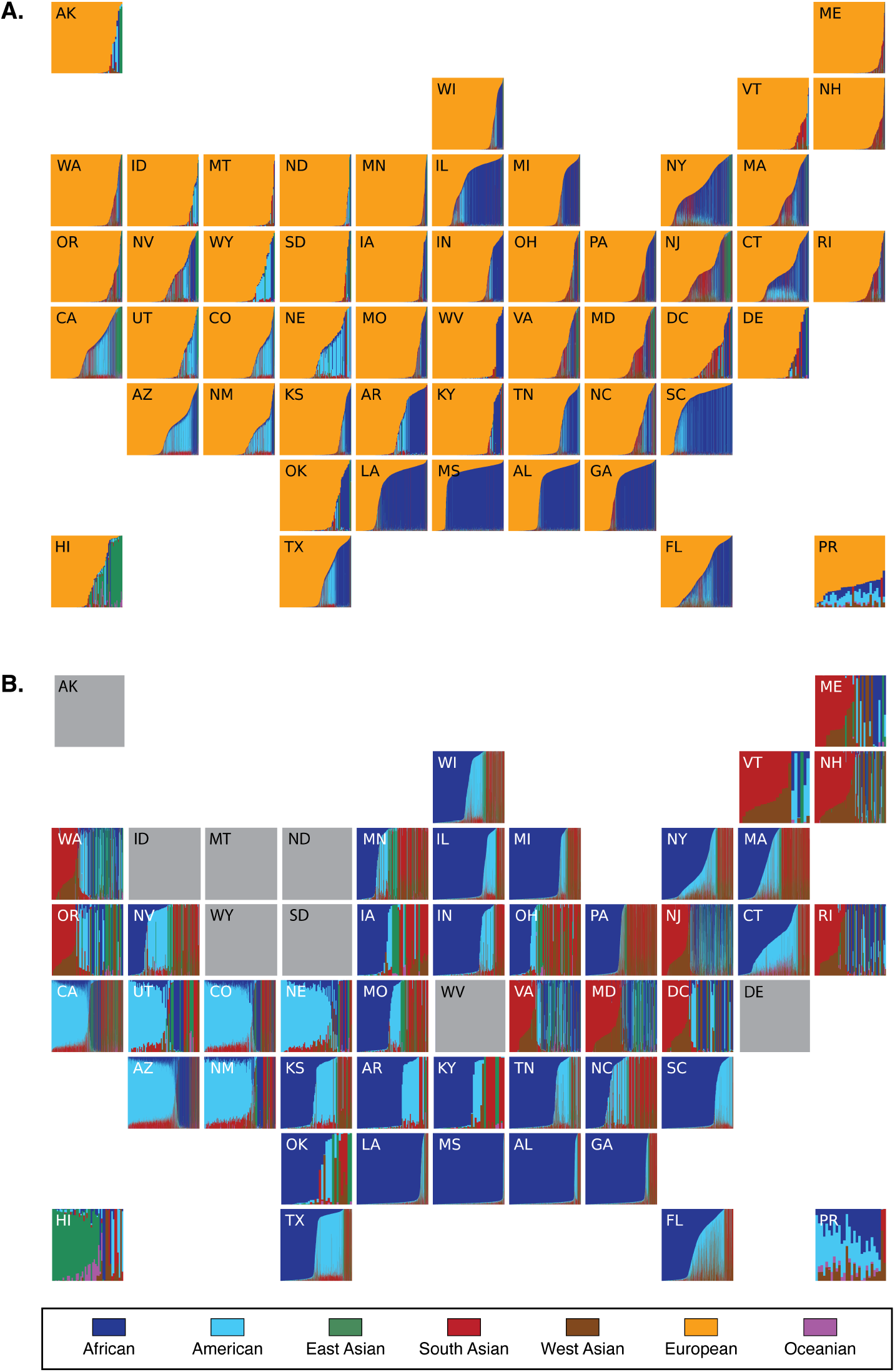
Genetic ancestry by geography. Genetic ancestry proportions are shown for *All of Us* participants sampled from the fifty US states and Puerto Rico. (A) All participants and ancestry components. (B) Non-European genetic ancestry proportions for all individuals with <90% European ancestry. The results for states shaded in grey are suppressed owing to <20 participants with <90% European ancestry.

The relationship between *All of Us* participants’ age and genetic ancestry was assessed using genetic admixture entropy, where higher values indicate a more diverse combination of ancestry components within individual genomes and lower values indicate more homogenous ancestry (Figure 5). Genetic admixture entropy is negatively correlated with participant age, indicating that younger participants have more diverse ancestry combinations than older participants.

**Figure 5.**
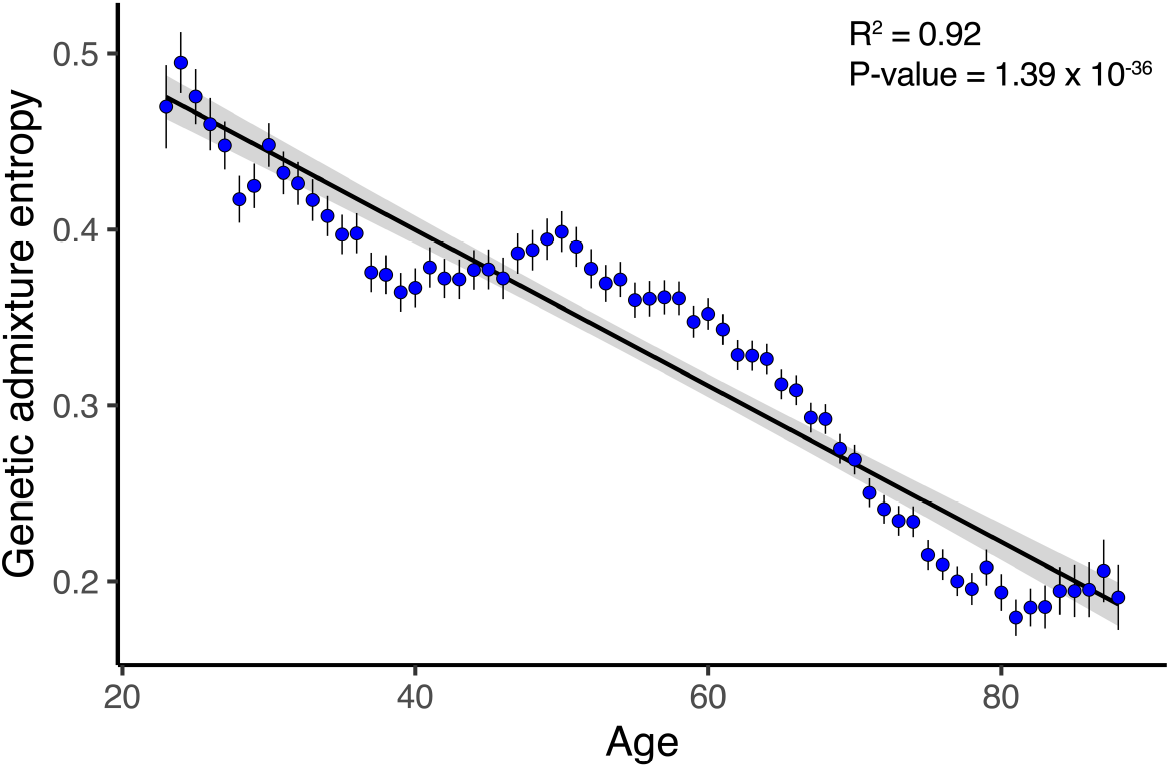
Genetic admixture by age. Genetic admixture entropy (y-axis) against participant age (x-axis). Ages shown in 100 bins with average and 95% CI values shown. Linear regression trend line shown with 95% CI shaded.

## Discussion

Our analysis demonstrates the genomic and ancestral diversity of the *All of Us* cohort, consistent with the project’s goals to recruit participants from population groups that are underrepresented in biomedical research in support of health equity. Indeed, *All of Us* is one of the most diverse population biomedical datasets in the world, and this represents an important step towards making precision medicine more widely available and more applicable to diverse communities in the US^7, 8, 40^. The promise of population biomedical datasets like *All of Us* rests on the integration of genetic, social, environmental, and health outcome data for many thousands of diverse participants. Given that genetic ancestry is derived from the genome, it should be possible to use genetic ancestry inference, together with population biomedical datasets, to help elucidate genetic and socioenvironmental contributions to health outcomes and disparities.

One challenge is that current methods for genetic ancestry inference, while accurate, are slow and do not scale to biobank sized datasets like *All of Us*. We developed the Rye algorithm as a fast and computationally efficient genetic ancestry inference method that can sale to biobank sized genomic data sets^33^. Application of Rye to genome-wide genetic data for 297,549 *All of Us* participants underscores its utility for this purpose. Using Rye, we found the *All of Us* cohort to be ancestrally diverse with distinct patterns of genetic ancestry and admixture among genetic similarity groups and geographic regions (Figures 2-4). The geographic patterns of genetic ancestry seen for the *All of Us* cohort are consistent with previous studies and could also reflect differences in participant recruitment across the country^41, 42, 43^.

The extent to which human genetic diversity is characterized by clusters of closely related individuals, i.e. population structure, versus clines of continuous genetic variation has long been a subject of interest^44, 45, 46, 47, 48^. The *All of Us* cohort allows for an assessment of the extent of population structure in the US given the large size of the cohort, the extensive sampling of participants across the country, and the demographic diversity of the participants. The application of several different cluster analysis methods to participants’ genomic PCA data revealed evidence for substantial population structure in the cohort, with dense clusters of relatively closely related participants interspersed among less dense regions in PC space (Figure 1). The population structure and genetic clusters that can be gleaned from clustering analysis of genomic PCA data are not readily apparent from visual inspection of these same data, owing to large size of the cohort and over-plotting of participants in dense regions of PC space (Figure 2A).

Finally, we show that genetic diversity in the US is increasing over time. Younger *All of Us* participants are far more ancestrally diverse than older participants, and this trend is evident across the entire age range of the cohort. This finding suggests that genetic ancestry categories and group designations will become increasingly obsolete over time^49^.

## Acknowledgements

We thank our colleagues, Kelsey Mayo, Ashley Able, Ashley Green, Andrea Ramirez, Anji Musick and Sokny Lim for providing their support and input throughout the demonstration project lifecycle. We thank Jennifer Zhang for providing input on the project’s code review. We thank Lee Lichtenstein and Jennifer Zhang for providing the data artifacts used for the project. We thank the DRC’s Research Support team for their help during implementation. We also thank the All of Us Science Committee and All of Us Steering Committee for their efforts in evaluating and finalizing the approved demonstration projects. The All of Us Research Program would not be possible without the partnership of contributions made by its participants. To learn more about the All of Us Research Program’s research data repository, please visit https://www.researchallofus.org/.

The All of Us Research Program is supported by the National Institutes of Health, Office of the Director: Regional Medical Centers: 1 OT2 OD026549; 1 OT2 OD026554; 1 OT2 OD026557; 1 OT2 OD026556; 1 OT2 OD026550; 1 OT2 OD 026552; 1 OT2 OD026548; 1 OT2 OD026551; 1 OT2 OD026555; IAA#: AOD 16037; Federally Qualified Health Centers: 75N98019F01202.; Data and Research Center: 1 OT2 OD35404; Biobank: 1 U24 OD023121; The Participant Center: U24 OD023176; Participant Technology Systems Center: 1 OT2 OD030043; Community Partners: 1 OT2 OD025277; 3 OT2 OD025315; 1 OT2 OD025337; 1 OT2 OD025276. In addition, the All of Us Research Program would not be possible without the partnership of its participants.

## SEEC Investigators

Priscilla E. Pemu, MD, MS^1^. Robert Meller DPhil, BSc^1^. Alexander Quarshie, MD^1^, MS. Kelley Carroll, MD^1^. Lawrence L. Sanders, MD^1^. Howard Mosby, CPA, CGMA^1^. Elizabeth I. Olorundare, MD, MPH^1^. Atuarra McCaslin, BS^1^. Chadrick Anderson, MHA. Andrea Pearson^1^. Kelechi C. Igwe, OD, MPH^1^. Karunamuni Silva^1^. Gwen Daugett, PMP. Jason McCray. Michael Prude. Chery^l^ Franklin, MD, MPH, FACOG^1^. Stephan Zuchner, M.D. Ph.D^2^. Olveen Carrasquillo, M.D. ^2^,. Rosario Isasi, JD. ^2^, MPH. Jacob L. McCauley, PhD^2^. Jose G Melo, MSPH^2^. Ana K Riccio, M.D. ^2^. Patrice Whitehead, MB (ASCP) ^2^. Patricia Guzman, MS^2^. Christina Gladfelter ^2^. Rebecca Velez, MA. ^2^, Mario Saporta, MD. ^2^ Brandon Apagüeño^2^, MS. Lisa Abreu, MPH^2^. Betsy Shenkman^3^. Bill Hogan^3^. Eileen Handberg^3^. Jamie Hensley^3^. Sonya White^3^. Brittney Roth-Manning^3^. Tona Mendoza^3^. Alex Loiacono^3^. Donny Weinbrenner^3^. Mahmoud Enani^3^. Ali Nouina^3^. Michael E. Zwick, Ph.D.^4^, Tracie C. Rosser, Ph.D.^4^, Arshed A. Quyyumi, M.D.^4^, Theodore M. Johnson II, M.D., MPH^4^, Greg S. Martin, M.D., M.Sc^4^, Alvaro Alonso, M.D., Ph.D.^4^, Tina-Ann Kerr Thompson, M.D^4^, Nita Deshpande, Ph.D. ^4^, H. Richard Johnston, Ph.D. ^4^, Hina Ahmed, MPH^4^, Letheshia Husbands, MPH^4^.

## Affiliations

1. Morehouse School of Medicine, Atlanta, Georgia, United States

2. University of Miami, Coral Gables, Florida, United States

3. University of Florida, Gainesville, FL

4. Emory University, Atlanta, GA.

## References

1. Bustamante CD, Burchard EG, De la Vega FM. Genomics for the world. Nature 475, 163–165 (2011).

2. Petrovski S, Goldstein DB. Unequal representation of genetic variation across ancestry groups creates healthcare inequality in the application of precision medicine. Genome Biol 17, 157 (2016).

3. Popejoy AB, Fullerton SM. Genomics is failing on diversity. Nature 538, 161–164 (2016).

4. Martin AR, Kanai M, Kamatani Y, Okada Y, Neale BM, Daly MJ. Clinical use of current polygenic risk scores may exacerbate health disparities. Nat Genet 51, 584–591 (2019).

5. All of Us Research Program I, et al. The “All of Us” Research Program. N Engl J Med 381, 668–676 (2019).

6. Ramirez AH, et al. The All of Us Research Program: Data quality, utility, and diversity. Patterns (N Y) 3, 100570 (2022).

7. Bianchi DW, et al. The All of Us Research Program is an opportunity to enhance the diversity of US biomedical research. Nat Med 30, 330–333 (2024).

8. Kathiresan N, Cho SMJ, Bhattacharya R, Truong B, Hornsby W, Natarajan P. Representation of race and ethnicity in the contemporary US health cohort All of Us Research Program. JAMA Cardiol 8, 859–864 (2023).

9. All of Us Research Program Genomics I. Genomic data in the All of Us Research Program. Nature 627, 340–346 (2024).

10. Pritchard JK. An Owner’s Guide to the Human Genome: an introduction to human population genetics, variation and disease.) (2023).

11. Hellenthal G, et al. A genetic atlas of human admixture history. Science 343, 747–751 (2014).

12. Nielsen R, Akey JM, Jakobsson M, Pritchard JK, Tishkoff S, Willerslev E. Tracing the peopling of the world through genomics. Nature 541, 302–310 (2017).

13. Royal CD, et al. Inferring genetic ancestry: opportunities, challenges, and implications. Am J Hum Genet 86, 661–673 (2010).

14. Wohns AW, et al. A unified genealogy of modern and ancient genomes. Science 375, eabi8264 (2022).

15. Mathieson I, Scally A. What is ancestry? PLoS Genet 16, e1008624 (2020).

16. Coop G. Genetic similarity versus genetic ancestry groups as sample descriptors in human genetics}. arXiv 2207.11595, (2023).

17. National Academies of Sciences Engineering and Medicine. Using Population Descriptors in Genetics and Genomics Research: A New Framework for an Evolving Field. The National Academies Press (2023).

18. Alexander DH, Novembre J, Lange K. Fast model-based estimation of ancestry in unrelated individuals. Genome Res 19, 1655–1664 (2009).

19. Maples BK, Gravel S, Kenny EE, Bustamante CD. RFMix: a discriminative modeling approach for rapid and robust local-ancestry inference. Am J Hum Genet 93, 278–288 (2013).

20. Bergstrom A, et al. Insights into human genetic variation and population history from 929 diverse genomes. Science 367, (2020).

21. Genomes Project C, et al. A global reference for human genetic variation. Nature 526, 68–74 (2015).

22. Chang CC, Chow CC, Tellier LC, Vattikuti S, Purcell SM, Lee JJ. Second-generation PLINK: rising to the challenge of larger and richer datasets. Gigascience 4, 7 (2015).

23. Conley AB, et al. A Comparative Analysis of Genetic Ancestry and Admixture in the Colombian Populations of Choco and Medellin. G3 (Bethesda) 7, 3435–3447 (2017).

24. Jordan IK, Rishishwar L, Conley AB. Native American admixture recapitulates population-specific migration and settlement of the continental United States. PLoS Genet 15, e1008225 (2019).

25. Nagar SD, et al. Genetic ancestry and ethnic identity in Ecuador. HGG Adv 2, 100050 (2021).

26. Hopkins B, Skellam JG. A new method for determining the type of distribution of plant individuals. Annals of Botany 18, 213–227 (1954).

27. Beygelzimer A, Kakadet S, Langford J, Arya S, Mount D, Li S. FNN: Fast nearest neighbor search algorithms and applications. (2024).

28. Venables WN, Ripley BD. Modern Applied Statistics with S, Fourth edn. Springer (2002).

29. McInnes L, Healy J, Astels S. hdbscan: Hierarchical density based clustering. J Open Source Softw 2, 205 (2017).

30. Manichaikul A, Mychaleckyj JC, Rich SS, Daly K, Sale M, Chen WM. Robust relationship inference in genome-wide association studies. Bioinformatics 26, 2867–2873 (2010).

31. Pedregosa F, et al. Scikit-learn: Machine learning in Python. Journal of machine Learning research 12, 2825–2830 (2011).

32. Patin E, et al. Dispersals and genetic adaptation of Bantu-speaking populations in Africa and North America. Science 356, 543–546 (2017).

33. Conley AB, et al. Rye: genetic ancestry inference at biobank scale. Nucleic Acids Res 51, e44 (2023).

34. All of Us Research Program. Genomic Research Data Quality Report.) (2022).

35. Bivand R, Keitt T, Rowlingson B. rgdal: Bindings for the ‘Geospatial’ Data Abstraction Library. (2023).

36. Hafen R. geofacet: ‘ggplot2’ Faceting Utilities for Geographical Data. (2023).

37. Medina-Rivas MA, et al. Choco, Colombia: a hotspot of human biodiversity. Rev Biodivers Neotrop 6, 45–54 (2016).

38. Homburger JR, et al. Genomic insights into the ancestry and demographic history of South America. PLoS Genet 11, e1005602 (2015).

39. Ruiz-Linares A, et al. Admixture in Latin America: geographic structure, phenotypic diversity and self-perception of ancestry based on 7,342 individuals. PLoS Genet 10, e1004572 (2014).

40. Abul-Husn NS, Kenny EE. Personalized medicine and the power of electronic health records. Cell 177, 58–69 (2019).

41. Dai CL, et al. Population histories of the United States revealed through fine-scale migration and haplotype analysis. Am J Hum Genet 106, 371–388 (2020).

42. Han E, et al. Clustering of 770,000 genomes reveals post-colonial population structure of North America. Nat Commun 8, 14238 (2017).

43. Bryc K, Durand EY, Macpherson JM, Reich D, Mountain JL. The genetic ancestry of African Americans, Latinos, and European Americans across the United States. Am J Hum Genet 96, 37–53 (2015).

44. Serre D, Paabo S. Evidence for gradients of human genetic diversity within and among continents. Genome Res 14, 1679–1685 (2004).

45. Rosenberg NA, et al. Genetic structure of human populations. Science 298, 2381–2385 (2002).

46. Rosenberg NA, Mahajan S, Ramachandran S, Zhao C, Pritchard JK, Feldman MW. Clines, clusters, and the effect of study design on the inference of human population structure. PLoS Genet 1, e70 (2005).

47. Mountain JL, Cavalli-Sforza LL. Multilocus genotypes, a tree of individuals, and human evolutionary history. Am J Hum Genet 61, 705–718 (1997).

48. Bowcock AM, Ruiz-Linares A, Tomfohrde J, Minch E, Kidd JR, Cavalli-Sforza LL. High resolution of human evolutionary trees with polymorphic microsatellites. Nature 368, 455–457 (1994).

49. Lewis ACF, et al. Getting genetic ancestry right for science and society. Science 376, 250–252 (2022).

