## Supplemental Information for "Genetic ancestry and population structure in the All of Us Research Program cohort"

Supplementary Figure 1. **Analysis flowchart.** The number of participants retained at each step are shown.

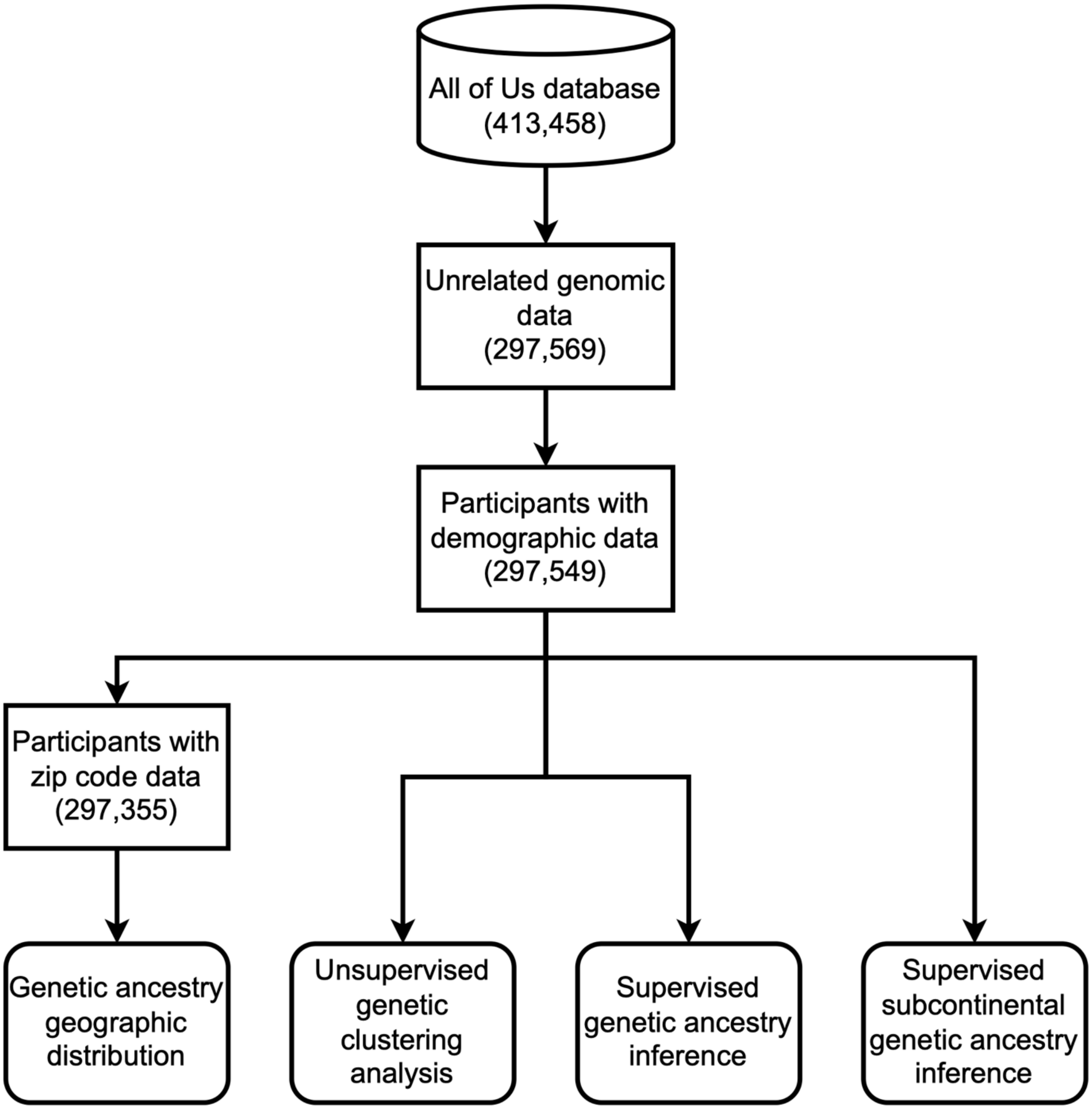

Supplementary Table 1. **Continental ancestry reference populations.**

| Population | Source[a] | Number of samples | Continental Ancestry Group |
|---|---|---|---|
| Balochi | HGDP | 17 | South Asian |
| Bedouin | HGDP | 15 | West Asian |
| Bougainville | HGDP | 9 | Oceanian |
| Brahui | HGDP | 20 | South Asian |
| CHB | 1KGP | 102 | East Asian |
| Colombian | HGDP | 6 | American |
| Dai | HGDP | 9 | East Asian |
| Druze | HGDP | 1 | West Asian |
| ESN | 1KGP | 99 | African |
| FIN | 1KGP | 85 | European |
| GBR | 1KGP | 87 | European |
| GIH | 1KGP | 2 | South Asian |
| GWD | 1KGP | 106 | African |
| IBS | 1KGP | 86 | European |
| ITU | 1KGP | 73 | South Asian |
| JPT | 1KGP | 104 | East Asian |
| Karitiana | HGDP | 12 | American |
| KHV | 1KGP | 96 | East Asian |
| LWK | 1KGP | 97 | African |
| Makrani | HGDP | 12 | South Asian |
| Maya | HGDP | 13 | American |
| MSL | 1KGP | 85 | African |
| Palestinian | HGDP | 41 | West Asian |
| Papuan Highlands | HGDP | 9 | Oceanian |
| Papuan Sepik | HGDP | 7 | Oceanian |
| PEL | 1KGP | 14 | American |
| Pima | HGDP | 12 | American |
| She | HGDP | 10 | East Asian |
| STU | 1KGP | 89 | South Asian |
| Surui | HGDP | 8 | American |
| TSI | 1KGP | 100 | European |
| Tujia | HGDP | 9 | East Asian |
| Tuscan | HGDP | 8 | European |
| YRI | 1KGP | 107 | African |

[a] 1KGP – 1000 Genomes Project[1], HGDP – Human Genome Diversity Project[2]

Supplementary Table 2. **Subcontinental ancestry reference populations.**

| Population | Source[a] | Number of samples | Subcontinental Ancestry Group |
|---|---|---|---|
| **African subcontinental ancestry** | | | |
| Ahizi | Patin et al | 20 | West Africa |
| Akele | Patin et al | 38 | Southwest Africa |
| Badwee | Patin et al | 38 | Southwest Africa |
| Baka | Patin et al | 108 | Rain Forest Hunter Gatherer |
| Bakota | Patin et al | 50 | Southwest Africa |
| BantuKenya | HGDP | 9 | Bantu |
| Bapunu | Patin et al | 49 | Southwest Africa |
| Bariba | Patin et al | 18 | West central Africa |
| Bateke | Patin et al | 44 | Southwest Africa |
| Bekwil | Patin et al | 5 | Southwest Africa |
| Benga | Patin et al | 40 | Southwest Africa |
| Biaka | HGDP | 22 | Rain Forest Hunter Gatherer |
| Duma | Patin et al | 40 | Southwest Africa |
| Eshira | Patin et al | 40 | Southwest Africa |
| ESN | 1KGP | 98 | West central Africa |
| Eviya | Patin et al | 20 | Southwest Africa |
| Fang | Patin et al | 67 | Southwest Africa |
| Fon | Patin et al | 12 | West central Africa |
| Galoa | Patin et al | 43 | Southwest Africa |
| GBR | 1KGP | 87 | European |
| GWD | 1KGP | 106 | West Africa |
| Kimbundu | Patin et al | 16 | Southwest Africa |
| Kongo | Patin et al | 9 | Southwest Africa |
| LWK | 1KGP | 93 | Bantu |
| Makina | Patin et al | 41 | Southwest Africa |
| MSL | 1KGP | 85 | West Africa |
| Ndumu | Patin et al | 36 | Southwest Africa |
| Nzebi | Patin et al | 62 | Southwest Africa |
| Obamba | Patin et al | 46 | Southwest Africa |
| Okande | Patin et al | 7 | Southwest Africa |
| Orungu | Patin et al | 19 | Southwest Africa |
| Ovimbundu | Patin et al | 11 | Southwest Africa |
| Shake | Patin et al | 49 | Southwest Africa |
| Tsogo | Patin et al | 60 | Southwest Africa |
| Umbundo | Patin et al | 5 | Southwest Africa |
| Yacouba | Patin et al | 17 | West Africa |
| YRI | 1KGP | 107 | West central Africa |
| **East Asian subcontinental ancestry** | | | |
| Austronesian | *All of Us* Inferred | 50 | Austronesian |
| CHB | 1KGP | 44 | Northern Han |

| | | | |
|---|---|---|---|
| CHS | 1KGP | 82 | Southern Han |
| CDX | 1KGP | 91 | Southeast Asian |
| Dai | HGDP | 9 | Southeast Asian |
| Han | HGDP | 4 | Northern Han |
| Japanese | HGDP | 27 | Japanese |
| JPT | 1KGP | 101 | Japanese |
| KHV | 1KGP | 84 | Southeast Asian |
| Korean | *All of Us* Inferred | 50 | Korean |
| **South Asian subcontinental ancestry** | | | |
| Balochi | HGDP | 17 | North Indian |
| Brahui | HGDP | 20 | North Indian |
| GIH | 1KGP | 2 | South Indian |
| ITU | 1KGP | 73 | South Indian |
| Kalash | HGDP | 22 | Central Asian |
| Makrani | HGDP | 12 | North Indian |
| STU | 1KGP | 89 | South Indian |
| **European subcontinental ancestry** | | | |
| FIN | 1KGP | 85 | Northern Europe |
| GBR | 1KGP | 87 | Northwestern Europe |
| IBS | 1KGP | 86 | Iberian Europe |
| TSI | 1KGP | 100 | Italian Europe |
| Tuscan | HGDP | 8 | Italian Europe |

[a] 1KGP – 1000 Genomes Project[1], HGDP – Human Genome Diversity Project[2], Patin et al.[3], *All of Us* Inferred – refers to dense PCA clusters of East Asian samples (identified with HDBSCAN) with locations in PCA space that are consistent with previous studies of Austronesian (i.e. Filipino) and Korean populations[4, 5].

Supplementary Methods. **Uniform Manifold Approximation and Projection (UMAP) analysis.** Uniform Manifold Approximation and Projection (UMAP) unsupervised dimension reduction and clustering was performed on the first 25 PCs using umap R package with 100 training epochs and a spread value of 229. Density-based clustering of genomic PCA and UMAP data was performed using the HDBSCAN algorithm30. HDBSCAN was run on the first 5 PCs for the PCA data and for the two UMAP embeddings with parameters min_samples=2,000 and min_cluster_size=2,500. Cluster boundaries were visualized for PCA and UMAP plots using the ggforce R package.

Supplementary Figure 2. **Cluster concordance between PCA and UMAP unsupervised clustering methods.** (A) PCA shows 7 density-based clusters, and (B) UMAP shows 13 density-based clusters. Color-codes for participants from the 7 PCA clusters are superimposed on the UMAP clusters. (C) Correspondence between PCA and UMAP clusters as shown by the percent of PCA participants (y-axis) that map to each UMAP cluster (x-axis).

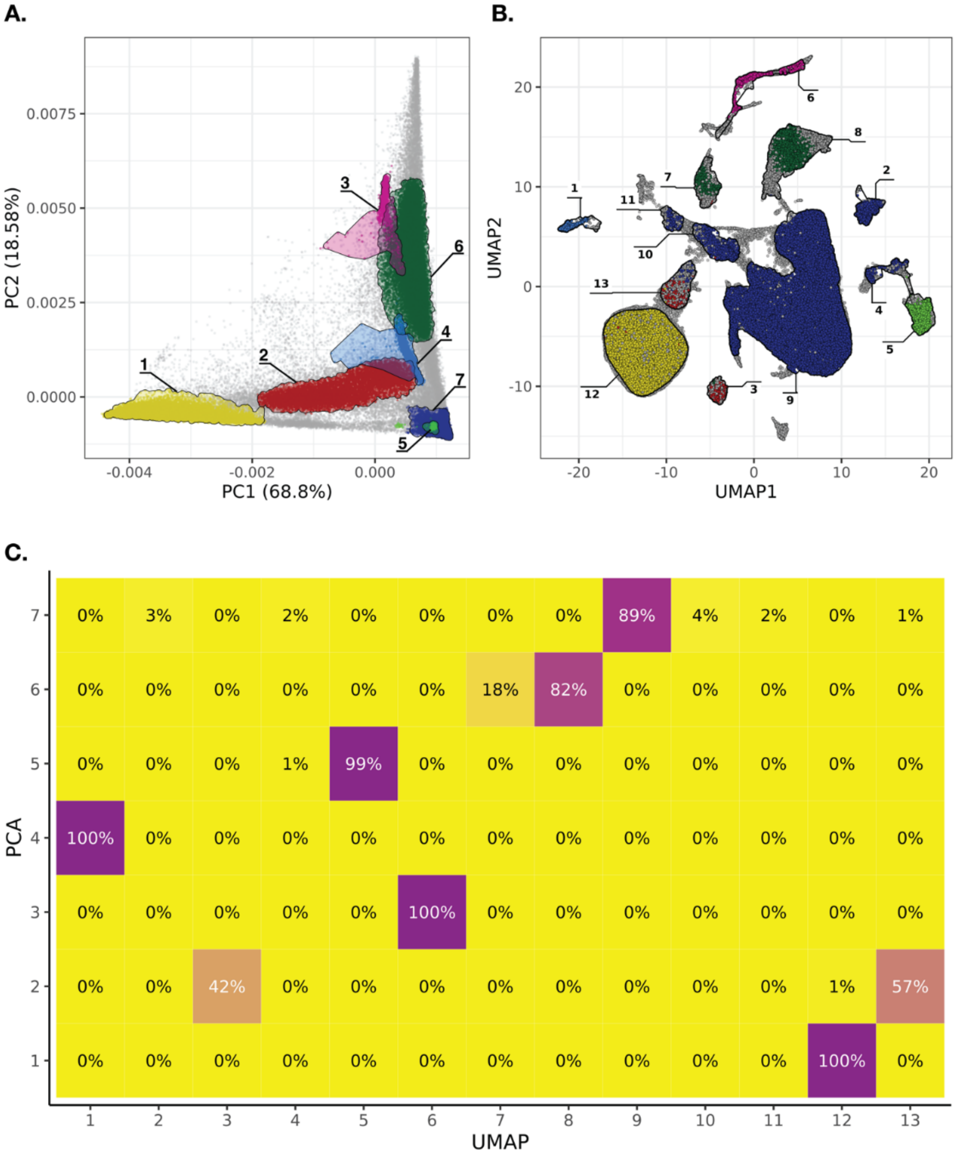

Supplementary Figure 3. **Genomic PCA for global reference populations.** (A) PC1 versus PC2 and (B) PC1 versus PC3. Percent variance explained by each PC are shown. Global reference population individuals are color-coded according to their geographic origins as shown in the key.

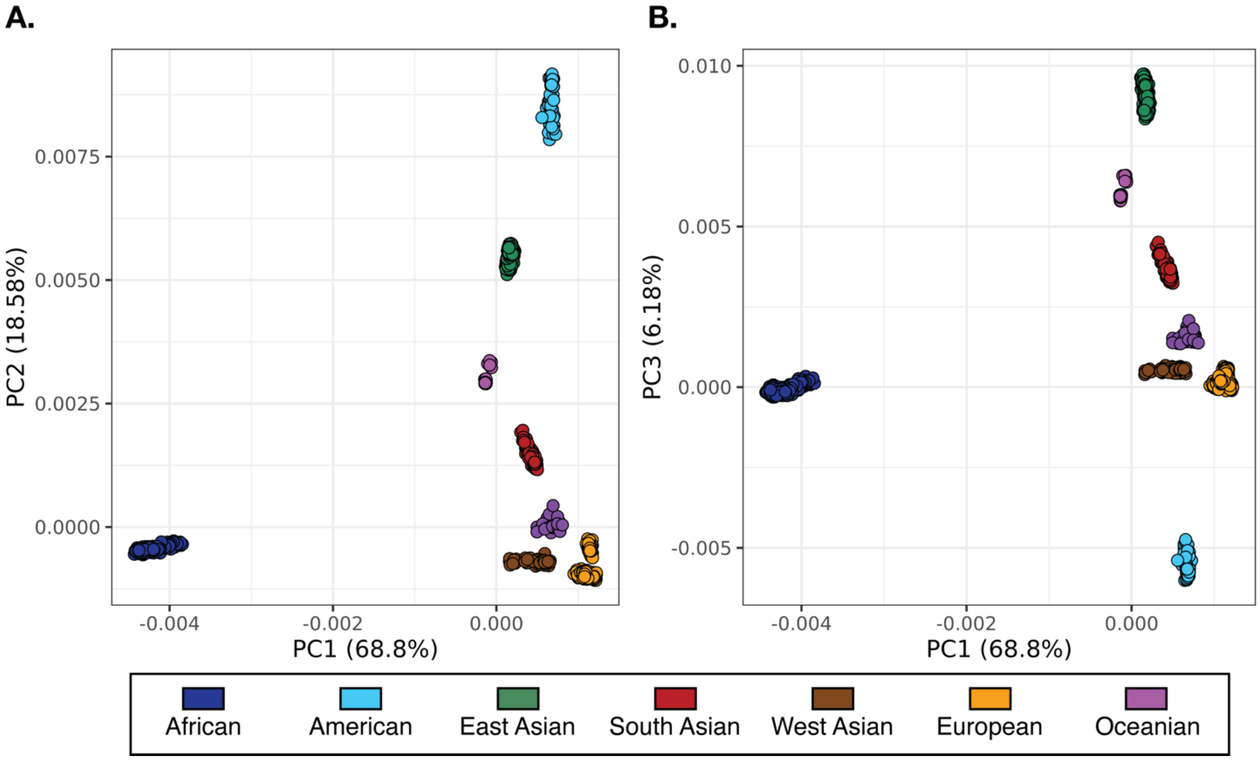

Supplementary Figure 4. **Predicted categorical continental ancestry versus inferred continuous genetic ancestry for *All of Us* participants.** The Rye algorithm used for this study infers continuous genetic ancestry percentages for *All of Us* participants. The *All of Us* Researcher Workbench provides categorical ancestry group predictions for participants. Average continental ancestry percentages (continuous genetic ancestry; x-axis) are shown for participants across predicted continental ancestry groups (categorical ancestry group; y-axis). Participants are assigned to continental ancestry groups if they are predicted to belong to a single group with >70% confidence. The Middle Eastern ancestry group provided by the Researcher Workbench corresponds to West Asian group described in this study.

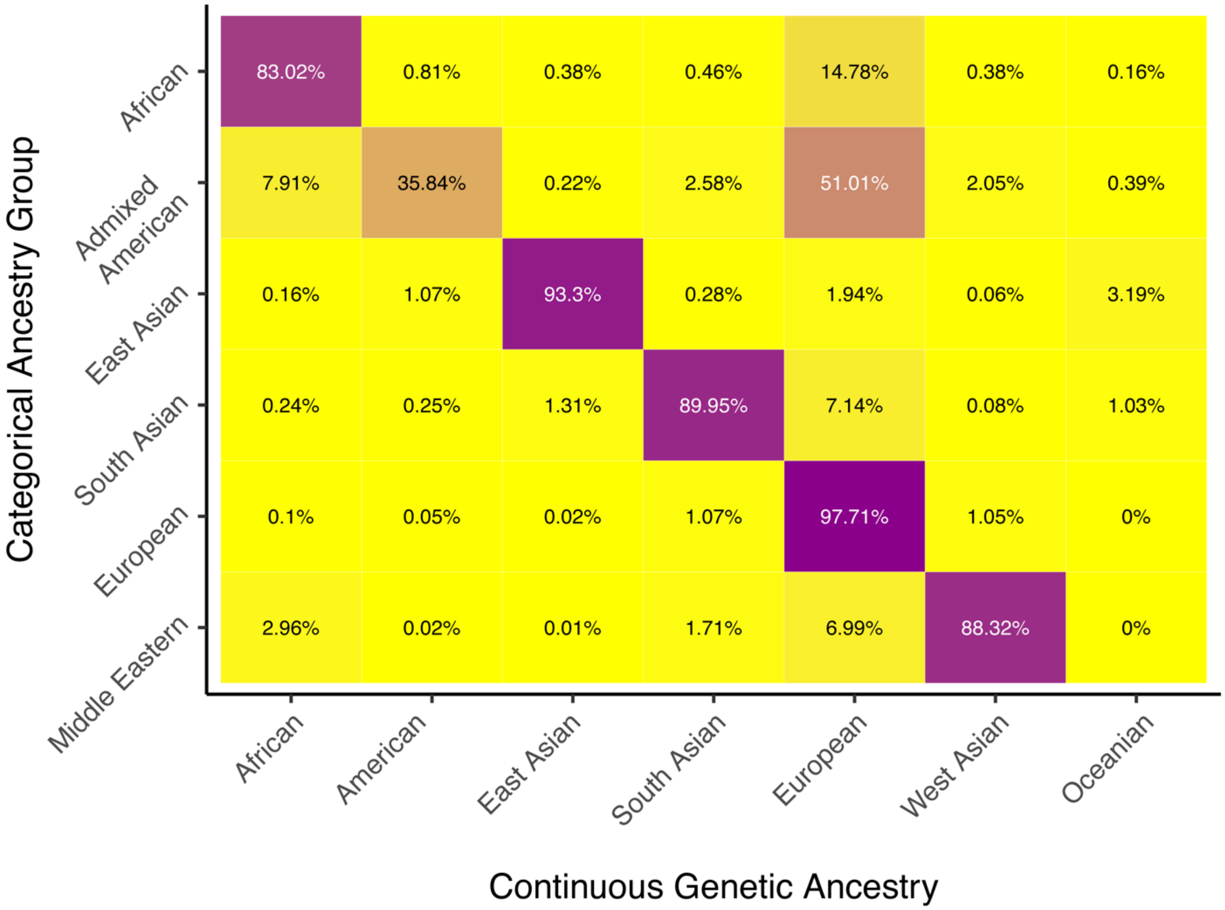

Supplementary Figure 5. **Coherent continental clusters used for subcontinental ancestry inference.** Genomic PCA with global reference populations are color-coded as shown, and yellow diamonds around each reference ancestry cluster (for African, East Asian, European, and South Asian ancestry) show the selected *All of Us* participants for whom subcontinental ancestry was inferred. (A) PC1 versus PC2 and (B) PC1 versus PC3.

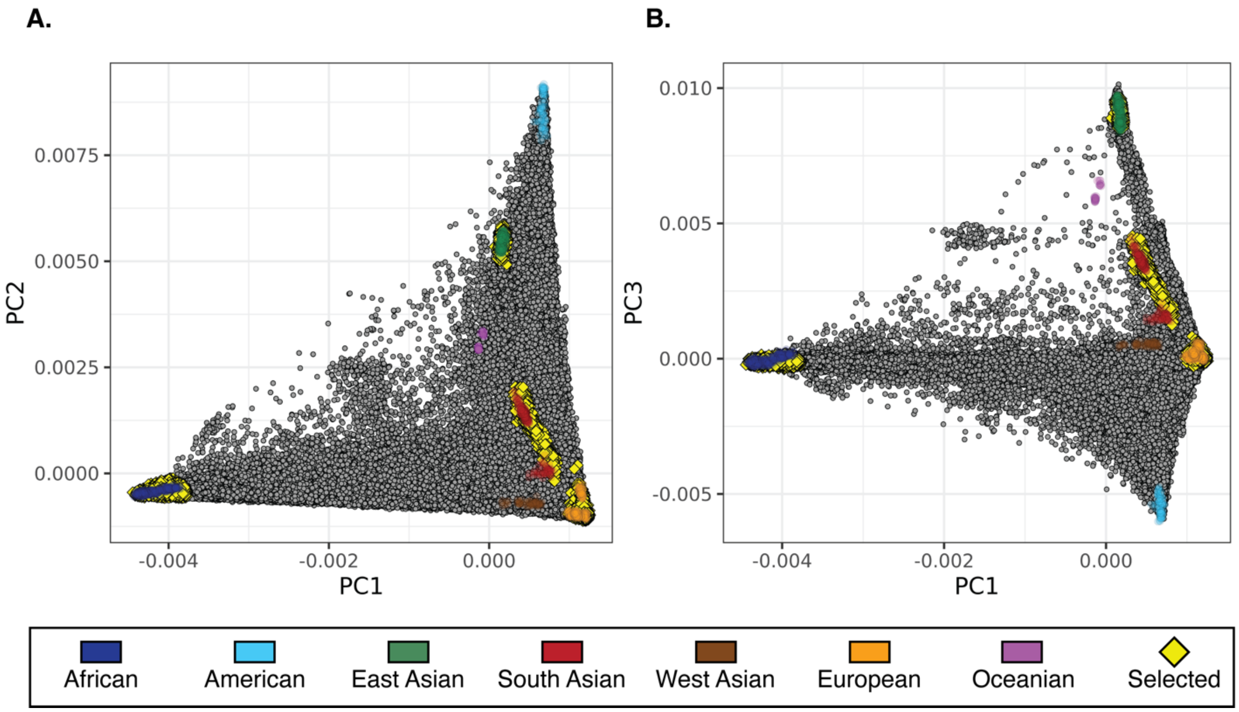

Supplementary Figure 6. **Genomic PCA for subcontinental reference populations.** *All of Us* participants are shown in gray. (A) African, (B) East Asian, (C) South Asian, and (D) European. (E) Continental ancestry percentages for the *All of Us* participants selected for subcontinental ancestry inference.

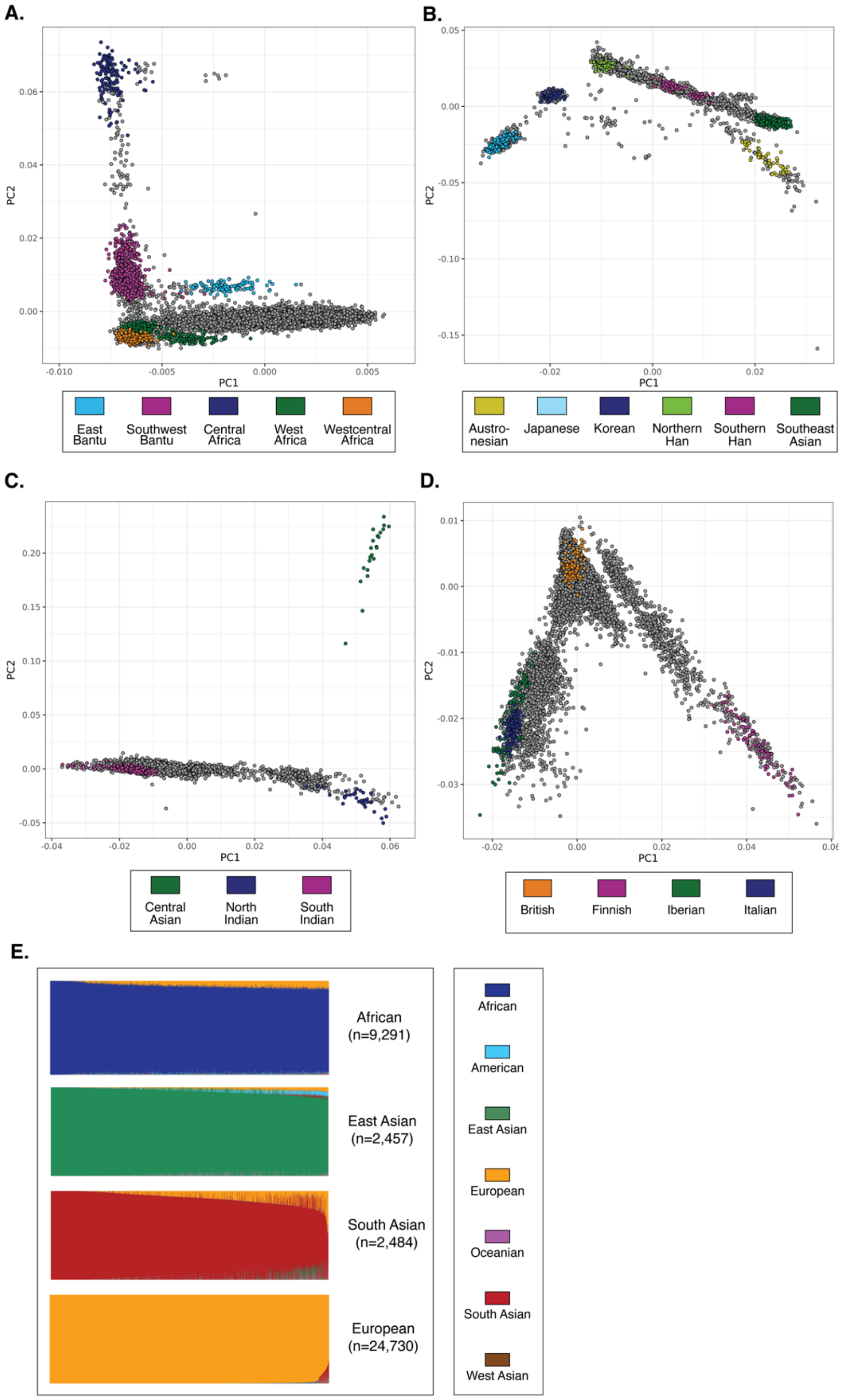

Supplementary Figure 7. **Genomic UMAP for subcontinental reference populations.** *All of Us* participants are shown in gray. (A) African, (B) East Asian, (C) South Asian, and (D) European continental ancestry groups.

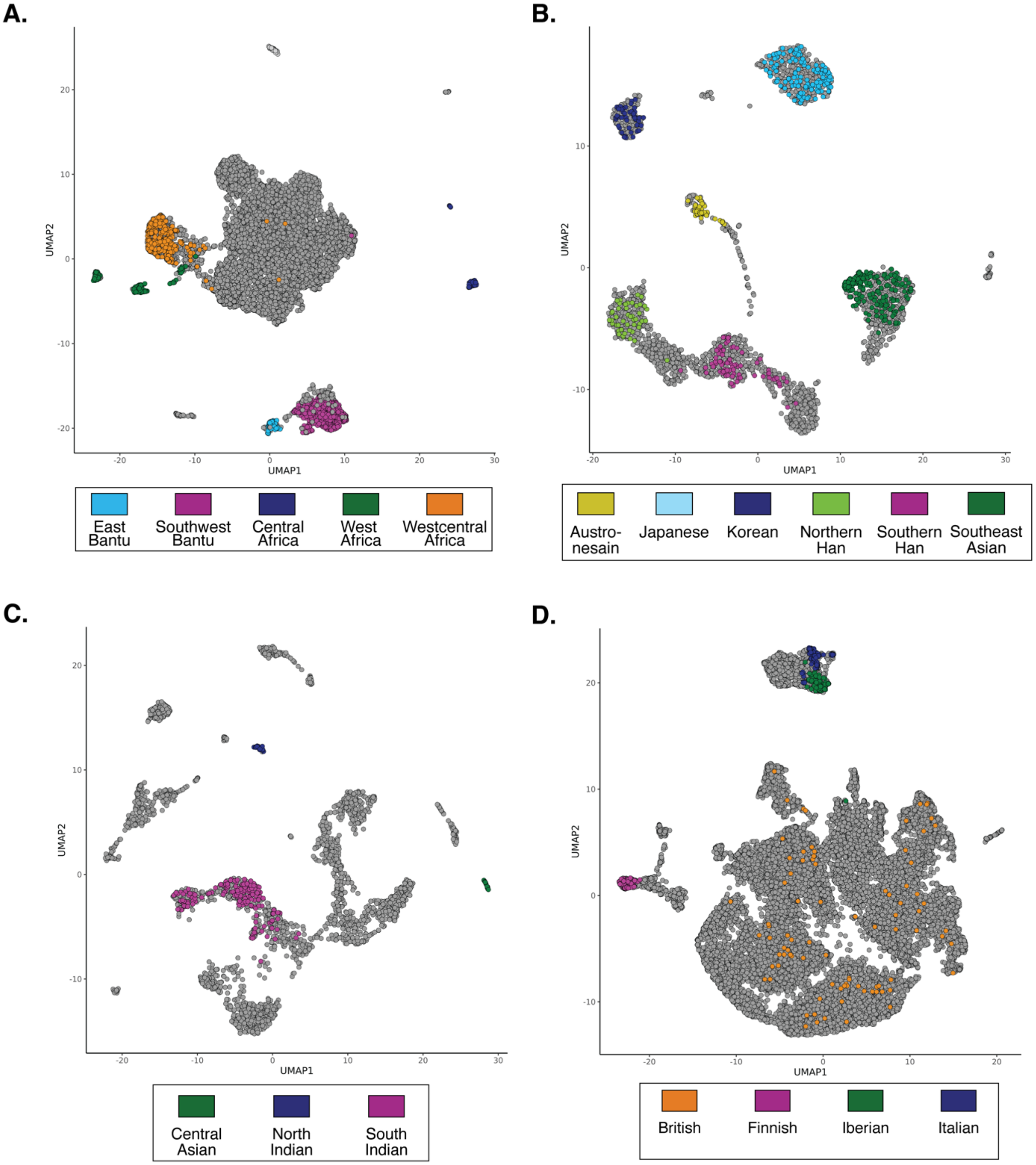

**Supplementary References**

1. Genomes Project C*, et al.* A global reference for human genetic variation. *Nature* **526**, 68-74 (2015).
2. Bergstrom A*, et al.* Insights into human genetic variation and population history from 929 diverse genomes. *Science* **367**,  (2020).
3. Patin E*, et al.* Dispersals and genetic adaptation of Bantu-speaking populations in Africa and North America. *Science* **356**, 543-546 (2017).
4. Zhang P*, et al.* NyuWa Genome resource: A deep whole-genome sequencing-based variation profile and reference panel for the Chinese population. *Cell Rep* **37**, 110017 (2021).
5. Tian C*, et al.* Analysis of East Asia genetic substructure using genome-wide SNP arrays. *PLoS One* **3**, e3862 (2008).
